# New Insights into How JUUL™ Electronic Cigarette Aerosols and Aerosol Constituents Affect SARS-CoV-2 Infection of Human Bronchial Epithelial Cells

**DOI:** 10.1101/2022.08.23.505031

**Authors:** Rattapol Phandthong, Man Wong, Ann Song, Teresa Martinez, Prue Talbot

**Affiliations:** Department of Molecular, Cell and System Biology, University of California, Riverside, CA 92521, USA

**Keywords:** E-cigarette, JUUL™, Nicotine, SARS-CoV-2, COVID-19, Air-liquid Interface, ACE2, TMPRSS2

## Abstract

**Background:** The relationship between the use of tobacco products and SARS-CoV-2 infection is poorly understood and controversial. Most studies have been done with tobacco cigarettes, while few have examined the effect of electronic cigarettes (ECs) on SARS-CoV-2 infection. We tested the hypothesis that EC fluids and aerosols with high concentrations of nicotine promote SARS-COV-2 infection by increasing viral entry into human respiratory epithelial cells.

**Methods:** Responses of BEAS-2B cells to authentic JUUL™ aerosols or their individual constituents (propylene glycol (PG)/vegetable glycerin (VG) and nicotine) were compared using three exposure platforms: submerged culture, air-liquid-interface (ALI) exposure in a cloud chamber, and ALI exposure in a Cultex® system, which produces authentic heated EC aerosols. SARS-CoV-2 infection machinery was assessed using immunohistochemistry and Western blotting. Specifically, the levels of the SARS-CoV-2 receptor ACE2 (angiotensin converting enzyme 2) and a spike modifying enzyme, TMPRSS2 (transmembrane serine protease 2), were evaluated. Following each exposure, lentivirus pseudoparticles with spike protein and a green-fluorescent reporter were used to test viral penetration and the susceptibility of BEAS-2B cells to infection.

**Results:** Nicotine, EC fluids, and authentic JUUL™ aerosols increased both ACE2 levels and TMPRSS2 activity, which in turn increased viral particle entry into cells. While most data were in good agreement across the three exposure platforms, cells were more responsive to treatments when exposed at the ALI in the Cultex system, even though the exposures were brief and intermittent. In the Cultex system, PG/VG, PG/VG/nicotine, and JUUL™ aerosols significantly increased infection above clean air controls. However, both the PG/VG and JUUL™ treatments were significantly lower than nicotine/PG/VG. PG/VG increased infection only in the Cultex® system, which produces heated aerosol.

**Conclusion:** Our data are consistent with the conclusion that authentic JUUL™ aerosols or their individual constituents (nicotine or PG/VG) increase SARS-CoV-2 infection. The strong effect produced by nicotine was modulated in authentic JUUL aerosols, demonstrating the importance of studying mixtures and aerosols from actual EC products. These data support the idea that vaping increases the likelihood of contracting COVID-19.

## Introduction

COVID-19 (corona virus disease-2019), a serious respiratory illness caused by the SARS-CoV-2 virus (severe acute respiratory syndrome coronavirus 2), has resulted in the death of over 1,000,000 people in the United States and an estimated 15,000,000 people worldwide [1, 2]. Smoking can lead to lung diseases, including cancer and increases in viral infection [3, 4], and while there has been intense interest in the relationship between tobacco product/nicotine use and COVID-19 infection, progression, and severity, this relationship is currently poorly understood and sometimes contradictory. Patient-derived data have sometimes led to the conclusion than smoking protects against COVID-19 [5, 6, 7], while most studies, including several recent meta-analyses, have concluded that smoking is a risk factor for the progression of COVID-19 [8, 9, 10] and that patients with a smoking history have a higher probability of developing more severe symptoms of COVID-19 [11].

Several studies have addressed the effect of smoking on COVID-19 experimentally using *in vitro* and animal models. SARS-CoV-2 viral fusion with the host cell is aided by TMPRSS2 (transmembrane serine protease 2), an enzyme that modifies the viral spike protein, enabling it to fuse with the host cell [12]. The ACE2 receptor (angiotensin converting enzyme 2) for SARS-CoV-2 spike protein is elevated in biopsies from the respiratory tract of human smokers compared to non-smokers, supporting the idea that smokers are more susceptible to COVID-19 [13, 14, 15, 16, 17]. Smith et al [13] also found that smokers had elevated cathepsin B, an enzyme involved in spike processing following infection. In a single cell meta-analysis across various tissues, smoking was correlated with increased levels of ACE2 and TMPRSS2, which may contribute to COVID-19 pathogenesis [18]. TMPRRS2 expression was differentially regulated in different respiratory cell types of smokers [18]. Studies that directly examined viral entry into human cells concluded that cigarette smoking increases SARS-CoV-2 infection [19, 20]. One group further showed that cigarette smoking inhibited the airway basal cell repair processes and reduced the response of the innate immune system by suppressing interferon β-1 [19], both factors that may contribute to the severity of the disease.

Electronic cigarettes (ECs) are nicotine delivery devices that have gained popularity in the last decade [21, 22, 23]. ECs heat e-liquids to produce aerosols that contain numerous chemicals, some of which are distinct from those in tobacco smoke [4, 24, 25]. EC aerosols can have multiple adverse effects on the respiratory system [22, 26, 27, 28, 29]. These include negative effects on lung physiology and the immune response, which may make fighting and recovery from a COVID-19 infection more difficult.

Several in vivo studies with mice and in vitro studies with human cells have investigated the effects of vaping and SARS-CoV-2 infection. Elevated ACE2 activities were found in the bronchial-lavage fluid from EC users and smokers [20]. Full body exposures to “Mint” but not “Mango” aerosols generated from JUUL™ pods increased ACE2 levels in mouse lung [30]. Three additional lung studies found elevated ACE2 levels following treatment with aerosols containing a mixture of PG (propylene glycol)/VG (vegetable glycerol) and nicotine [31,32, 33]. While these studies reported gender differences, they were not in agreement with respect to the gender affected.

The effects of EC use and ACE2 levels has been studied in vitro with cultured human cells from the respiratory system. BEAS-2B cells treated with e-liquids in submerged culture showed that PG/VG elevated ACE2 mRNA expression, and the effect was dependent on other chemicals in the e-liquid mixture [34]. JUUL™ “Virginia Tobacco” e-liquid increased infection in primary tracheobronchial and small airway epithelium in submerged culture [20]; however, e-liquid rather than authentic aerosol was tested.

The relationship between vaping authentic EC aerosols and acquisition of COVID-19 is poorly understood. To add clarity to this topic, we tested the hypothesis that EC aerosols increase SARS-CoV-2 infection by increasing levels of ACE2 in human bronchial epithelium, increasing levels and activities of TMPRRS2, and increasing viral infection. We performed controlled laboratory experiments with human cells that were exposed to known concentrations of both JUUL™ “Virginia Tobacco” EC fluids and aerosols, nicotine, PG and VG, and then examined discrete endpoints relevant to infection. We chose JUUL™ “Virginia Tobacco” because this product is currently marketed, popular, and has few flavor chemicals [35], making its fluid and aerosol a relatively simple mixture for testing. Moreover, tobacco flavored products are likely to remain marketable in the future as they are not subject to the FDA’s enforcement policy that proposes to remove flavored EC products from the market [36]. We tested both JUUL™ “Virginia Tobacco” fluid and aerosols equivalent to those inhaled during vaping. Testing was done across three *in vitro* exposure platforms: submerged cultures, cloud chamber ALI exposure, and Cultex® ALI exposure. Submerged culture has historical importance and is still frequently used [37, 38, 39, 40, 41]. The cloud chamber enables individual chemicals to be studied without using heat to create the aerosol, so that endpoint effects can be attributed to a particular chemical(s) without interference from other chemicals, solvents, or reaction products [42]. The Cultex® system was used to vape JUUL™ ECs by heating their fluid to create authentic JUUL™ aerosols, which are equivalent to those an EC user inhales. Cultex®-generated aerosols contain the chemicals normally present in the fluid [25] plus any reaction products or metals added when fluid is heated in an EC atomizer [43, 44, 45, 46, 47, 48, 49] Infection of cells was evaluated using SARS-CoV-2 pseudoparticles with a green-fluorescent reporter protein. Data were compared across exposure platforms and across exposure groups (JUUL™ fluids, JUUL™ aerosols, and individual chemicals).

## Materials and Methods

### BEAS-2B, HEK 293, and HEK 293T^ACE2^ Cell Culture and Maintenance

BEAS-2B cells (ATCC, Manassas, VA, USA) were cultured in Bronchial Epithelial Basal Medium (BEBM; Lonza, Walkersville, MD, USA) supplemented with the Bronchial Epithelial Growth Bullet Kit (BEGM; Lonza, Walkersville, MD, USA) without the gentamycin antibiotic. Cells were cultured in Nunc T-25 tissue culture flasks (Fisher Scientific, Tustin, CA, USA), pre-coated overnight with collagen type I, bovine serum albumin, and fibronectin. Cells were passaged at 80% confluency. Cell cultures were washed with Dulbecco’s phosphate buffered saline without calcium or magnesium (DPBS-; Lonza, Walkersville, MD, USA) and detached with 0.25% trypsin with 0.53 mM EDTA (ATCC, Manassas, VA, USA) and poly-vinyl-pyrrolidone (Sigma-Aldrich, St. Louis, MO, USA). Cells were seeded at 3,000 cells/cm^2^ in a pre-coated T-25 flask, and culture medium was replaced every other day.

For in vitro submerged treatments, cells were seeded at a density of 9,000/cells/cm^2^ in pre-coated 6-well and 12-well plates and allowed to attach overnight prior to treatments. For ALI culture, cells were seeded at a density of 12,000 cells/cm^2^ in pre-coated 12-well Transwell® inserts with a pore size of 0.4 μm (Corning, Inc., Corning, NY, USA) and allowed to form a monolayer. Once a monolayer formed, the medium on the apical side of the transwell was removed, and the monolayer was acclimated to air for 24 hrs prior to an exposure. Medium in the basal-lateral side of the transwell was replaced every other day. Cells were incubated at 37°C, 5% CO2, and 95% relative humidity.

HEK 293T cells (ATCC, Manassas, VA, USA), and HEK 293T cells over expressing ACE2 (HEK 293T^ACE2^, BEI resource, Manassas, VA, USA; NR-52511) were cultured in DMEM high glucose medium supplemented with 10% fetal bovine serum (FBS; ATCC Manassas, VA, USA). At 80-90% confluency, cells were washed and detached as described above. Cells were seeded at 3,000 cells/cm^2^ and incubated in a 37°C, 5% CO_2_, 95% relative humidity incubator.

### Submerged Treatments

Refill fluids contained PG (Thermofisher, Tustin CA, USA) and VG (Thermofisher, Tustin CA, USA). At 80% confluency, BEAS-2B cells were treated for 24 hours with nicotine, PG/VG, PV/VG with nicotine, or JUUL™ fluids. All treatments were diluted with BEGM culture medium to reach working concentrations. Liquid (-)- nicotine (Sigma-Aldrich, St. Louis, MO, USA) was diluted to reach final concentrations of 0.03 mg/mL or 0.3 mg/mL. PG/VG (30/70 ratio) was diluted to reach a final concentration of 0.5% in volume. Stock solutions of PG/VG with nicotine were made and then diluted with culture medium to 0.5% PG/VG with either 0.03 mg/mL nicotine or 0.3 mg/mL nicotine. JUUL™ “Virginia Tobacco” was diluted to 0.5%, which contains PG/VG at 0.5% concentration and nicotine at 0.3 mg/mL.

### Nicotine Aerosol Exposure at the ALI in the VITROCELL® Cloud Chamber

Monolayers of BEAS-2B cells cultured on 12 well Transwell® inserts were placed into a VITROCELL® cloud chamber (VITROCELL® Walkirch, Germany) for an ALI exposure of various chemical aerosols that were generated without heating. Prior to exposures, pre-warmed culture medium was dispensed into wells of the exposure chamber and allowed to equilibrate to 37 °C. Stock nicotine was diluted with PBS- to make exposure solutions with final concentrations of either 0.03 mg/mL or 0.3 mg/mL nicotine. For each exposure, 200 μL of exposure solution was added into a VITROCELL® nebulizer to generate a uniform aerosol with a flow rate of 200 μL/min without using heat. Control cells were exposed to PBS- aerosols.

BEAS-2B cells cultured on 12-well inserts were exposed to 1 puff of PBS- and nicotine (0.03 or 0.3 mg/mL) aerosol during a 1.5-minute aerosol generation period followed by a 3-minute aerosol deposition period. After the exposure, the cells were returned to the incubator to recover for 24 hrs.

### EC Aerosol Exposure at the ALI in the Cultex® RFS Compact Exposure System

A Cultex® RFS compact exposure module (Cultex® Laboratories GmbH, Hannover, Germany) was used to expose cells to humified sterile air (clean air control) or EC aerosols generated from EC devices. Prior to each exposure, cells were placed into the exposure chambers, which contained culture medium maintained at 37°C.

The Cultex® exposure system consisted of a sampling module and an aerosol guiding module. The sampling module contained a custom designed EC smoking robot (RTI International, North Carolina, USA) that draws filtered air from the biosafety cabinet or EC aerosol from the EC device into a 200 mL syringe. After collecting a 55 mL sample of either filtered air or EC aerosols, the Cultex® system then dispensed the sample into the aerosol guiding module, where it was combined with humified zero air (1 L/min). This mixture step diluted the sample and generated a uniform flow before the exposure mixtures were directly distributed onto each biological sample. Exposure mixtures were allowed to settle onto cells for 5 seconds, then vented out of the exposure chamber at a flow rate of 5 mL/min, generated by a mass flow controller (Boekhorst, Bethlehem, PA, USA), and finally dispensed into a waste container.

Each exposure consisted of 55 mL of filtered air or EC aerosol generated using a 4 second puff duration and a 30 second puff interval. BEAS-2B cells were exposed to 10 puffs of either clean air or EC aerosol and allowed to recover for 24 hrs prior to analyses.

### Quantification of Deposited Nicotine in the ALI Exposure Systems

To quantify the nicotine deposited in the ALI exposure systems, exposures were repeated as described in the previous protocol sections using isopropyl alcohol (IPA; Fisher Scientific, Fair Lawn, NJ, USA) in place of the biological samples. Ten mL of IPA were added to each exposure well in both the cloud chamber and Cultex® system. After the exposures, the IPA was collected and shipped overnight on ice to Portland State University where it was analyzed using gas chromatography-mass spectrometry (GC-MS). Chemical analysis was performed with an Agilent 5975 GC/MS system (Agilent, Santa Clara, CA, USA) using an internal standard-based calibration procedure and method previously described [25, 50].

### Immunocytochemistry of BEAS-2B cells

BEAS-2B cells were seeded in 8-well chamber slides (Ibidi, Gräfelfing, Germany). At 80% confluency, cells were treated with nicotine or EC fluids for 24 hrs in an incubator. Cells were then fixed in 4% paraformaldehyde for 15 mins at room temperature and washed several times with DPBS+. Samples were permeabilized with 0.1% Triton X and blocked using donkey serum (Sigma-Aldrich, St. Louis, MO, USA) for 1 hour at room temperature, followed by an overnight incubation in primary antibody. After several washes with PBS-T (DPBS+0.1% Tween), the samples were incubated at room temperature in the dark for 2 hrs with appropriate secondary antibodies. Samples were washed several times and mounted using Vectashield with DAPI (Vectashield, San Francisco, CA, USA). Fluorescent cells were imaged with a Nikon Eclipse Ti inverted microscope (Nikon Instrument, Melville, NY, USA) using a 60X objective, and images were captured using a high-resolution Andor Zyla VSC-04941 camera (Andor, Belfast, UK). Antibodies used were anti-ACE2 (1:100; R&D System, Minneapolis MN, USA), anti-TMPRSS2 (1:200; Santa Cruz, Dallas, TX, USA), Secondary antibodies were Alexa fluor-488 or Alexa fluor-594 (Thermofisher, Tustin CA, USA).

### Lysate Preparation for Western Blot and Proteolytic Assay

After submerged treatments or exposures at the ALI, RIPA buffer with or without PMSF protease inhibitors (ChemCruz Biochemical, Dallas, TX, USA) was used to lyse cells. Lysates for Western blots were prepared using RIPA buffer with protease inhibitors, while lysates for proteolytic assays used RIPA buffer without protease inhibitors. The cell lysates were vortexed every 10 mins for 30 mins, pipetted through 23-gauge needles several times, then centrifuged at 3,000 x g for 5 mins at 4°C. The lysate protein was quantified using the Pierce BCA assay kit (Thermo Scientific, Waltham, MA. USA). Each Western blot used 30 μg of protein, and each proteolytic assay used 5 μg of protein.

### Western Blotting

Following lysate preparation, denaturing buffer (β-mercaptoethanol and Laemeli buffer, 1:10) was added to each Western blot lysate at a 1:4 ratio. The buffer/lysate mixtures were heated at 95°C for 2 mins, then loaded onto an SDS gel (BioRad, Carlsbad, CA, USA) for electrophoretic separation of proteins (120V for 1-2 hrs), and afterwards transferred to a BioRad PVDF membrane at 200mA overnight at 4°C. Following transfer, the membrane was cut horizontally, either below or above the expected location of the protein of interest based on its molecular weight (kDa). The membranes were then blocked with 5% milk in TBS-T (TBS with 1%Tween-100) buffer for 2 hrs. and incubated overnight at 4°C with antibodies against ACE2 (1:400; R&D systems, Minneapolis, MN, USA), TMPRSS2 (1:1000; Santa Cruz, Dallas, TX, USA), and GADPH (1:2000; Cell Signaling Technology, Danvers, MA, USA). Next, the membranes were washed for 30 mins in TBS-T, then incubated in an HRP-conjugated secondary antibody (1:1000; Santa Cruz, Dallas, TX, USA or Cell Signaling Technology, Danvers, MA, USA) for 2 hrs. at room temperature. Finally, the membranes were developed using BioRad Clarity™ Western ECL Substrate reagent (BioRad, Carlsbad, CA, USA) in a BioRad ChemiDoc™ Imaging System (BioRad, Carlsbad, CA, USA), which determined the optimal exposure time for each protein (ACE2 = 120 ms; TMPRSS2 = 24 ms; GADPH = 4 ms).

### TMPRSS2 Proteolytic Assay

The TMPRSS2 enzyme assay was performed using a modification of a previously published protocol [51]. The fluorogenic substrate, Boc-Gln-Ala-Arg-AMC · HCl, (Bachem, Torrance, CA. USA) was dissolved in DMSO and diluted in reaction buffer (50 mM Tris pH 8, 150 mM NaCl) to a final concentration of 10 μm. The fluorogenic substrate was added to each well of a 96-well plate and the fluorescence intensity was measured at 340/440 nm over a 1 hr period at 37 °C using a Bio-Tek Synergy HTX (Agilent, Santa Clara, CA, USA) plate reader.

### Spike Viral Pseudoparticle Production

Pseudoparticle production was performed as outlined in Crawford et al. [52]. In brief, HEK293T cells were plated with antibiotic-free medium at a density of 7 x 10^6/^/T75 flask and transfected with Lipofectamine3000 (Thermo Fisher Sci, Waltham, MA. USA) using a total of 15 μg of BEI lentiviral plasmids (NR-52520: pHAGE2-CMV-ZsGreen-W, 7.5 μg; NR-52517: HDM-Hgpm2, 1.65 μg; NR-52519: pRC-CMV Rev1b, 1.65 μg; NR-52518: HDM-tat1b, 1.65 μg; NR-52314: pHDM-SARS-CoV-2 Spike, 2.55 μg), 60 μL of Lipofectamine, and 20 μL of P3000 reagent following the manufacturer’s protocol. After overnight incubation, fresh medium supplemented with 1% bovine serum albumin (BSA; Sigma-Aldrich, St. Louis, MO, USA) was added to the cells. Fluorescence microscopy was used to visually inspect transfection efficiency and the expression of ZsGreen. Wild-type HEK293T cells, which were not transfected by the lentiviral plasmids, were used as a transfection control. The cell culture medium from transfected and wild-type HEK293T cells was collected 48 hours after transfection, centrifuged, and the resulting supernatant was filtered using a 0.45 μm syringe filter. The filtered supernatant was mixed with 5X PEG (Abcam, Cambridge, UK) and precipitated overnight at 4°C. The lentivirus was collected by centrifugation and resuspension of the pellet in Viral Re-suspension Solution (Abcam, Cambridge, UK). The virus aliquots were stored at −80°C. Prior to infection experiments, the transduction efficiency of each viral batch was determined by infecting 293T^ACE2^ cells and quantifying the number of infected cells with flow cytometry. The medium from transfection control cells were processed the same, and produced a mock transfection solution, which did not contain virus and was used for mock infection.

### Viral Pseudoparticle Infection

For each viral pseudoparticle infection experiment, a 0.3 multiplicity of infection (MOI) was used to infect BEAS-2B cells. Viral pseudoparticles were delivered as a mixture with the appropriate fresh culture medium

In the submerged culture infection experiments, cells were treated for 24 hours, then the treatment was replaced with pseudoparticle medium. In the ALI culture infection experiments, following the recovery period, 100 μL of pseudoparticle medium was added directly onto the apical side of the Transwell®.

In all infection experiments, cells or tissues were incubated with viral pseudoparticles for 24 hrs. Subsequently, the pseudoparticles were removed and the cells or tissues were allowed to incubate another 24 hrs to amplify the expression of the green fluorescence reporter protein in the infected cells. Cells were harvested and analyzed with flow cytometry to determine the number of infected cells.

Prior to flow cytometry, fluorescence microscopy was used to validate the expression of ZsGreen signal. All samples were pipetted several times and passed through a 35 μm filter of a Falcon™ Round-Bottom 5 mL polystyrene test tube (Fisher Scientific, Tustin, CA, USA) to generate single cell suspensions. Forward-Scatter-Height (FSC-H) and Forward-Scatter-Area (FSC-A) were used to generate a gate to select single cells for each sample. A gate to exclude small debris was created using Side-Scatter-Area (SSC-A) and FSC-A. Non-fluorescent mock infected cells, which were incubated in viral re-suspension solution without viral pseudoparticles, were used to produce a gate to quantify the fluorescence signal of infected cells. Final results were represented as percent of fluorescent infected cells in each sample.

### Statistical Analysis

In all cases, three independent experiments were performed using different passages of BEAS-2B cells. Statistical analyses were done using Minitab Statistics Software (Minitab, State College, PA, USA). Data were first checked for normality of distribution and homogeneity of variances. When data did not satisfy the assumptions of analysis of variance (ANOVA), they were subjected to a Box-Cox transformation, and again checked to verify that data satisfied the ANOVA model. Statistical analyses were done using one-way ANOVA. When significance was found (p< 0.05), data were further analyzed using Tukey’s multiple comparison posthoc test to isolate significant differences to specific groups. Means were considered significantly different for p < 0.05. Data were plotted using GraphPad Prism 7 software (GraphPad, San Diego, CA, USA)

## Results

### Effect of JUUL™ Fluids, Aerosols, and EC Chemicals (Nicotine and PG/VG) on ACE2 in BEAS-2B Cells During Submerged Culture and ALI Exposures

The effects of JUUL™ “Virginia Tobacco” fluids and aerosols and individual components in JUUL™ products (specifically nicotine and PG/VG) on ACE2 levels were examined in BEAS-2B cells using submerged and ALI exposures (**Figure 1**). BEAS-2B cells were treated in submerged culture for 24 hours with 0.5% of 5% nicotine JUUL™ “Virginia Tobacco” fluid, which contained 0.3 mg/mL of nicotine after dilution, nicotine (0.03 mg/mL or 0.3 mg/mL), 0.5% PG/VG (ratio = 30/70), or 0.5% PG/VG with nicotine (0.03 mg/mL or 0.3 mg/mL) (**Figure 1A**). Micrographs of treated BEAS-2B cells labeled with ACE2 antibodies showed increased expression of ACE2 in those groups that contained 0.3 mg/mL of nicotine (**Figure 1B**). Increased expression was confirmed using Western blots (**Figures 1 C-D**). Nicotine increased ACE2 dose dependently, and the increase was significant in those groups that contained 0.3 mg/mL of nicotine. ACE2 also increased in the JUUL™ “Virginia Tobacco” treated group, but it was not significantly different than the control, even though JUUL™ has 0.3 mg/mL of nicotine. Overall, nicotine treatment for 24 hours in submerged cultures increased ACE2 levels dose dependently. PG/VG alone increased ACE2, but the effect was not significant.

**Figure 1.**
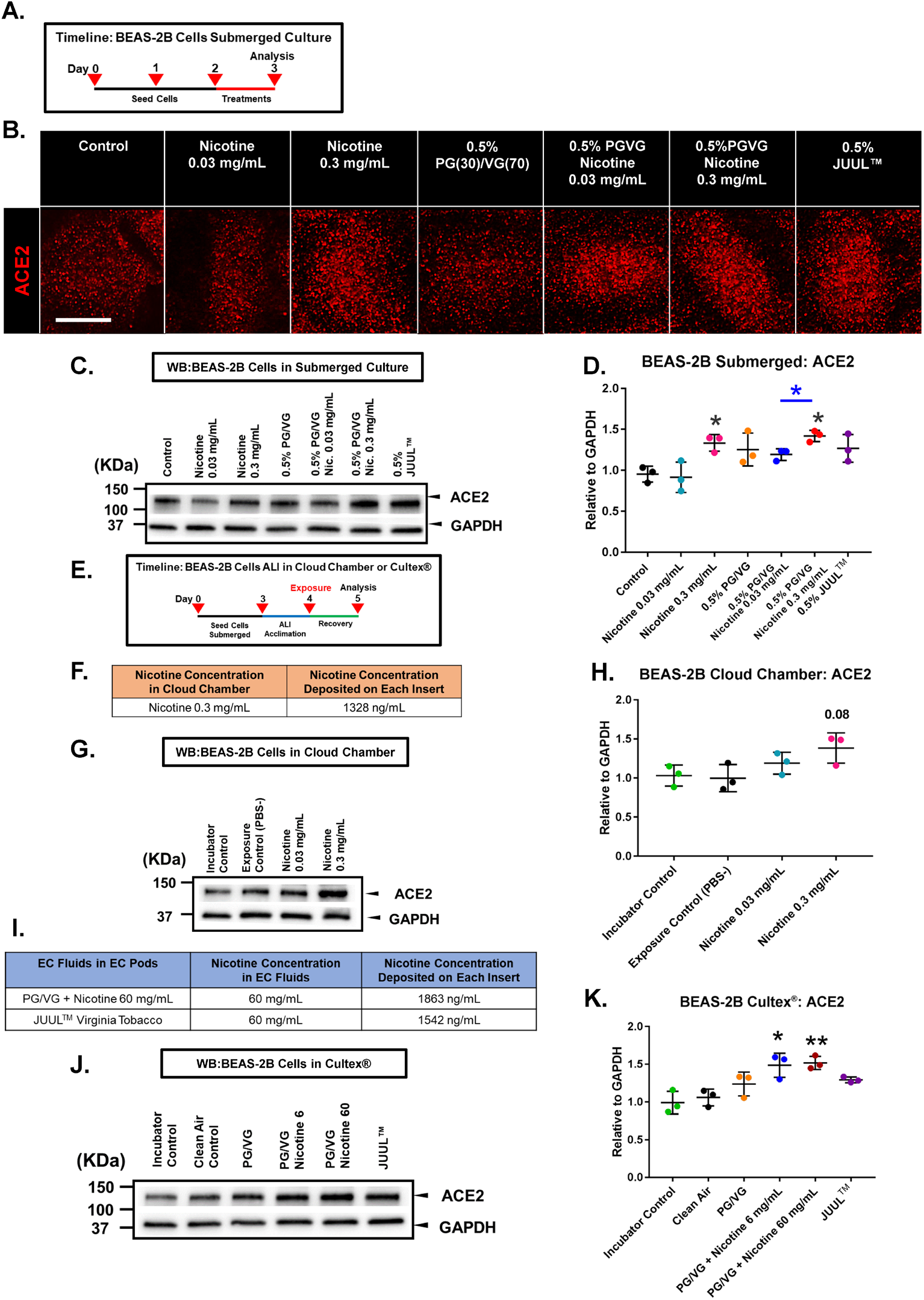
ACE2 Expression in BEAS-2B Cells During Exposure in Submerged Culture and at the Air-liquid Interface. **(A)** Experimental treatment used in submerged culture. **(B)** Micrographs showing cells labeled with ACE2 antibody. Scale bar = 50 μm. **(C - D)** The effect of JUUL™ e-liquid, nicotine, and PG/VG on ACE2 during submerged culture (western blot). **(E)** Experimental treatment used with cells exposed at the ALI in a cloud chamber or the Cultex® system. **(F)** Concentration of nicotine deposited in each well following aerosolization of 0.3 mg/mL of nicotine in a cloud chamber. **(G - H)** The effect of 1 puff of nicotine-containing aerosol on ACE2 following ALI exposure in a cloud chamber (western blot). **(I)** Nicotine concentration in each well following exposure to aerosol generated from PG/VG/nicotine 60 mg/mL or JUUL™ “Virginia Tobacco” in a Cultex® system. **(J - K)** The effect of 10 puffs of nicotine-containing EC aerosols on ACE2 following ALI exposure in a Cultex® system. The aerosols were generated from PG/VG, PG/VG with nicotine, and a JUUL™ EC. Data are plotted as the mean ± standard deviation of three independent experiments. Box-Cox transformed data were analyzed using a one-way ANOVA followed by Tukey’s test to compare means. Black * = significantly different than the control. Color * = significantly different groups. *p < 0.05, **p < 0.01.

Similar experiments were performed in which BEAS-2B cells were exposed to aerosols at the ALI in a VITROCELL® cloud chamber or Cultex® exposure system (**Figures 1 G-K**). In the cloud chamber, BEAS-2B cells were exposed to either 1 puff of aerosol generated from solutions of PBS or nicotine (0.03 mg/mL or 0.3 mg/mL) in PBS, then allowed to recover for 24 hours before Western blotting (**Figures 1G-H**). ACE2 increased in both nicotine-treated groups and was close to significance (p = 0.08) in the 0.3 mg/mL treated cells (**Figure 1H**). There was no significant difference between the incubator and PBS controls, indicating that incubation in the cloud chamber did not affect the cells.

In the Cultex® system, BEAS-2B cells were exposed to either 10 puffs of humidified, filtered clean air or 10 puffs of aerosols made using an authentic JUUL™ “Virginia Tobacco” EC. In addition, the JUUL™ EC battery was used with refillable third-party pods containing lab made refill fluids to determine which components in JUUL™ fluid affected ACE2 levels. The fillable pods contained PG/VG, PG/VG with 6 mg/mL of nicotine, or PG/VG with 60 mg/mL of nicotine. There was no significant difference between the incubator and clean air controls. PG/VG and authentic JUUL™ “Virginia Tobacco” aerosols elevated ACE2 expression, but not significantly (**Figure 1J and K**). Aerosols from **l**ab made fluids containing PG/VG and nicotine increased ACE2 expression significantly compared to the clean air control.

In both submerged and ALI exposures, nicotine and EC aerosols with nicotine resulted in a concentration-dependent elevation of ACE2 levels in BEAS-2B cells.

The exposures in the cloud chamber and Cultex® experiments were compared by measuring nicotine deposited in the transwell culture medium after exposure. When aerosolized in the cloud chamber, solutions containing 0.3 mg/mL of nicotine deposited 1,328 ng/mL of nicotine into the fluid in each insert (**Figure 1F**). In the Cultex® system, 1,863 ng/mL of nicotine were deposited into each insert from aerosols generated using PG/VG with 60 mg/mL of nicotine. JUUL™ aerosols deposited a similar concentration of nicotine into the fluid in each insert (1,543 ng/mL) (**Figure 1I**). These results demonstrate that similar amounts of nicotine were deposited in each insert in the two ALI exposure systems.

### The Effect of JUUL™ Fluids, PG/VG, and Nicotine on the Concentrations and Activities of TMPRSS2 in BEAS-2B Cells During Submerged Exposure

Submerged treatments were done to determine the effects of PG/VG, nicotine, and JUUL™ “Virginia Tobacco” fluid on TMPRSS2 levels and enzymatic activity (**Figure 2**). TMPRSS2 is critical for infection, as it modifies SARS-CoV-2 spike protein after binding to ACE2 and facilitates virus and host membrane fusion.

**Figure 2.**
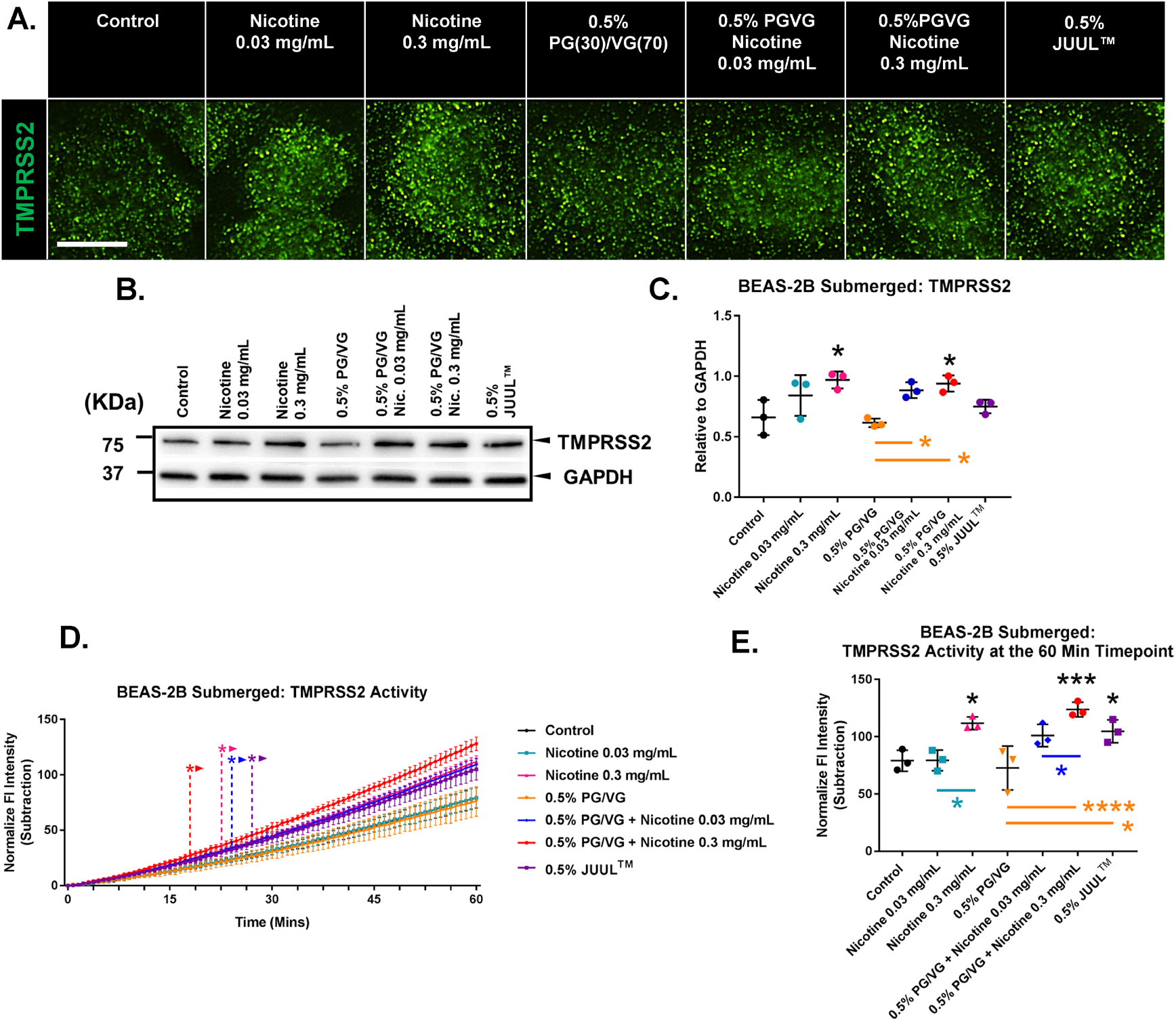
Effect of JUUL™ E-liquid, Nicotine, and PG/VG on TMPRSS2 in BEAS-2B Cells in Submerged Culture. **(A)** Micrographs showing cells labeled with TMPRSS2 antibody. Scale bar = 50 μm. **(B - C)** Effect of JUUL™ e-liquid, nicotine, and PG/VG on TMPRSS2 levels in submerged culture (western blot). **(D - E)** TMPRSS2 activity was increased by JUUL™ e-liquid and nicotine following 24 hours of treatment in submerged cultures. (**D**) Changes in activity over 60 minutes. **\*** Indicate when values became significantly different than the control (*p* ranged from 0.05 to 0.0001). **D** was analyzed using a two-way ANOVA followed by Dunnett’s compared to the control. (**E**) Activity at the 60-minute timepoint. All graphs show the mean ± standard deviation of three independent experiments. In **C** and **E** Box-Cox transformed data were analyzed using a one-way ANOVA followed by Tukey’s test. Black * = significantly different than the control. Color * = significantly different groups. * = p < 0.05, ** = p < 0.01, *** = p < 0.001, **** = p < 0.0001.

Similar exposure experiments were done to determine if TMPRSS2 is affected by JUUL™ “Virginia Tobacco” fluid, PG/VG, or nicotine. Treated BEAS-2B cells labeled with TMPRSS2 antibodies showed increased expression of TMPRSS2 in those groups exposed to 0.3 mg/mL of nicotine (**Figure 2A**). In Western blots, TMPRSS2 expression was not significantly affected by JUUL™ fluid or PG/VG alone (**Figure 2 B, C**). However, groups treated with lab-made fluids containing nicotine had elevated levels of TMPRSS2, an effect that was dose dependent (**Figure 2B-C**). TMPRSS2 activity, as measured by cleavage of a specific fluorescent substrate, increased (p = 0.07) when BEAS-2B cells were exposed to authentic JUUL™ “Virginia Tobacco” EC fluid. TMPPRS2 activity was dose dependently increased by nicotine and was significantly greater than the control in groups having 0.3 mg/mL of nicotine. TMPRSS2 activity was not significantly affected by PG/VG alone (**Figures 2D, 2E**).

### The Effect of Authentic JUUL™ Aerosols, PG/VG, and Nicotine Aerosols on TMPRSS2 Concentration and Activity in BEAS-2B Cells Following ALI Exposures

Similar experiments were performed with BEAS-2B cells exposed at the ALI to aerosols generated in a cloud chamber or Cultex® exposure system. TMPRSS2 levels were evaluated in Western blots, and its activity was analyzed using specific fluorogenic substrates.

In the cloud chamber, BEAS-2B cells were exposed to 1 puff of aerosol generated from solutions of either PBS or nicotine (0.03 mg/mL or 0.3 mg/mL) in PBS, then allowed to recover for 24 hours before Western blotting and analysis of proteolytic activities (**Figures 3A-D)**. Nicotine exposures did not significantly affect TMPRSS2 levels (**Figures 3A, 3B)**. However, the enzymatic activity was significantly increased in 0.3 mg/mL of nicotine (**Figure 3C, 3D**).

**Figure 3.**
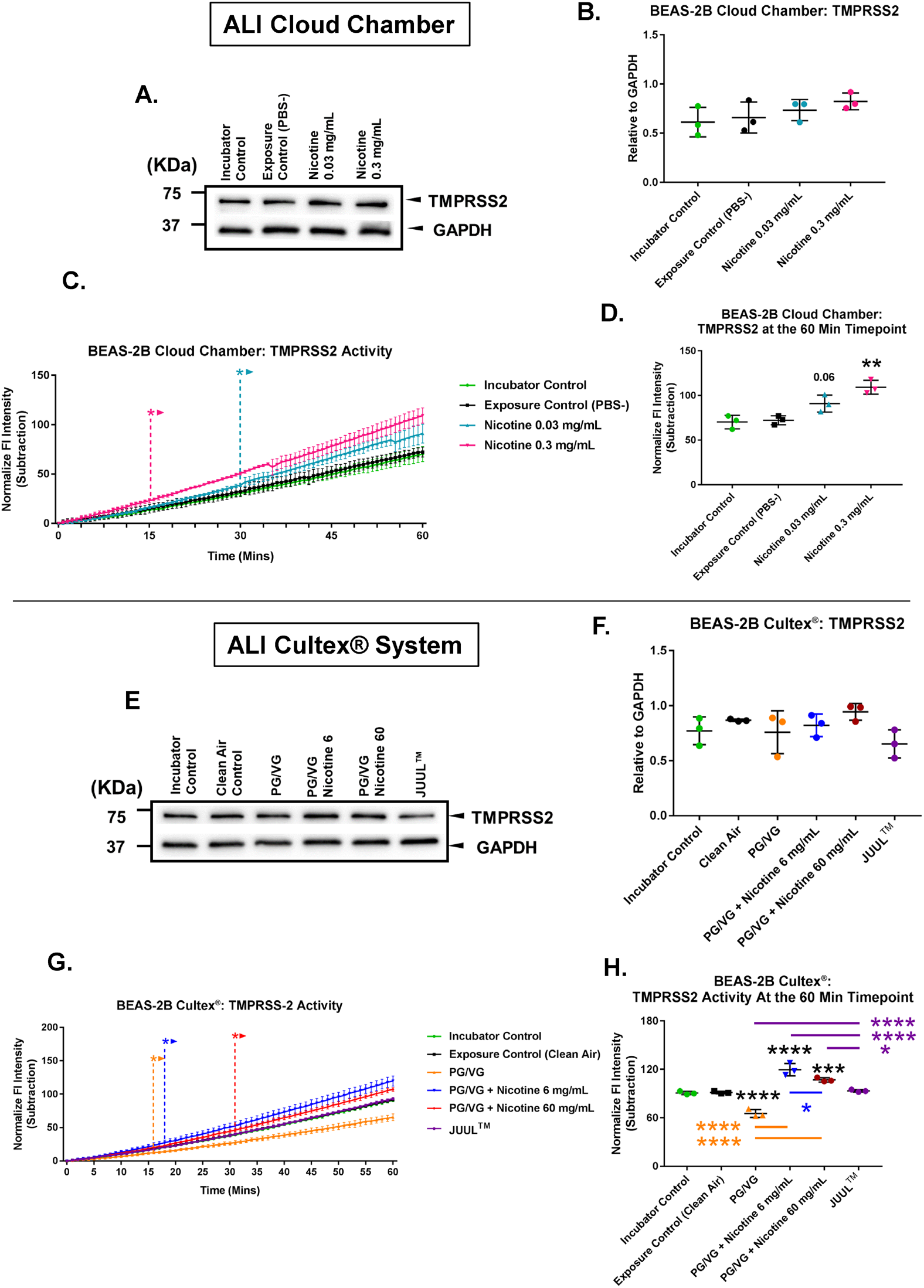
Effect of EC Aerosols on TMPRSS2 Activity in BEAS-2B Cells Exposed at the Air-liquid Interface. **(A - B)** Effect of 1 puff of pure nicotine aerosol on TMPRSS2 levels; exposure was at the ALI in a cloud chamber (Western blot). **(C - D)** TMPRSS2 activity was increased by nicotine following exposure at the ALI in a cloud chamber. (**C**) Changes in activity over 60 minutes. **\*** Indicate when values became significantly different than the control (*p* ranged from 0.05 to 0.0001). (**D**) Activity at the 60-minute timepoint. **(E - F)** Western blot showing the effect of 10 puffs of PG/VG, PG/VG/nicotine, and JUUL™ EC aerosol on TMPRSS2 levels during ALI exposure in the Cultex® system. **(G - H)** Nicotine increased TMPRSS2 activity during ALI exposure in the Cultex® system. PG/VG decreased activity. JUUL™ aerosols increased activity (**G**) Changes in activity over 60 minutes. * Indicate when values became significantly different than the control (*p* ranged from 0.05 to 0.0001). (**H**) Activity at the 60-minute timepoint. All graphs are plotted as means ± standard deviation of three independent experiments. Following a Box-Cox transformation, **B, D, F** and **H** were analyzed using a one-way ANOVA followed by Tukey’s test to compare means. Black * = significantly different than the control. Color * = significantly different groups. **C** and **G** were analyzed using a two-way ANOVA followed by Dunnett’s test to compare means to the control. *p < 0.05, **p < 0.01, ***p < 0.001, ****p < 0.0001.

In the Cultex® system, BEAS-2B cells were exposed to 10 puffs of aerosol containing JUUL™ “Virginia Tobacco” aerosol, PG/VG only, or PG/VG with nicotine (**Figure 3E-3H**). TMPRSS2 levels did not differ significantly in any of the treatments when compared to the clean air control group (**3E, 3F**). However, TMPRSS2 activity decreased significantly in the PG/VG group, increased significantly in the two groups containing PG/VG with nicotine, but did not change significantly in the authentic JUUL™ aerosol group, which was generated from fluid containing 60 mg/L of nicotine. (**Figure 3G, 3H**).

These results demonstrate that aerosols containing nicotine plus PG/VG can elevate TMPRSS2 activity in BEAS-2B cells, which may facilitate infection by enabling more rapid cleavage of the viral spike protein after ACE2 binds to host cells.

### JUUL™ Fluids and Aerosols and Nicotine Increased Viral Pseudoparticle Infection of BEAS-2B Cells in Submerged Cultures and ALI Exposures

Experiments were done to determine how authentic JUUL™ fluids and aerosols, and their components (nicotine and PG/VG) affect infection of BEAS-2B cells. SARS-CoV-2 viral pseudoparticles were constructed using lentivirus as the infecting agent. The pseudoparticles contain SARS-CoV-2 spike protein in their envelope and a reporter plasmid that encodes ZsGreen, which is used to identify infected cells.

293T cells, genetically modified by stable transfection to overexpress ACE2 (293T^AEC2^), were used to test viral pseudoparticle infectivity, determine pseudoparticle density, and optimize the expression of ZsGreen. 293T^ACE2^ cells were infected using an MOI of 0.3 (**Figure 4A**). These data were used to optimize infection of BEAS-2B cells (**Figure 4B-4H).**

**Figure 4.**
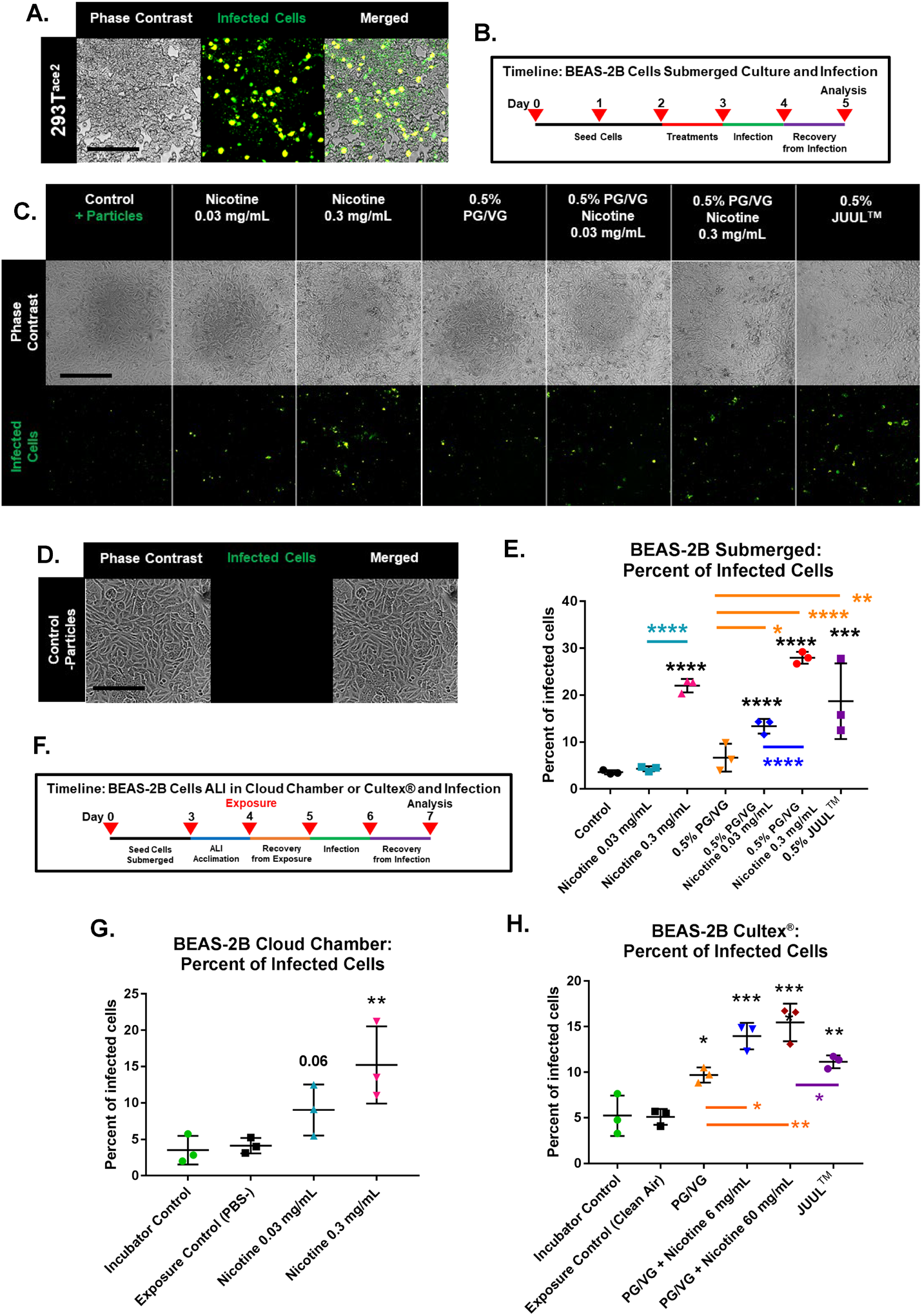
JUUL™ Aerosols and Nicotine Increased Infection After Submerged Treatment or ALI Exposure to Aerosols. **(A)** Micrographs showing 293T^ace2^ cells infected by SARS-CoV-2 viral pseudoparticles. Infected cells express ZsGreen. Scale bar = 250 μm. **(B)** Experimental treatments in submerged culture followed by infection. **(C - E)** Micrographs and flow cytometry showing infection of cells treated 24-hrs with EC chemicals or JUUL™ fluid then infected for 24 hrs. (**C - D**) Micrographs showing (**C**) cells infected with pseudoparticles and (**D**) uninfected cells. (**E**) Percent of infected cells determined by flow cytometry. Scale bars in C and D = 250 μm. **(F)** Experimental treatment used with cells exposed at the ALI in a cloud chamber or a Cultex® system, then infected with viral pseudoparticles for 24 hours. **(G)** Percent of infected cells after cloud chamber exposure to 1 puff of PBS (control), 0.03 mg/mL nicotine, or 0.3 mg/mL nicotine, followed by 24 hrs of exposure to pseudoparticles. **(H)** Percent of infected cells after exposure to 10 puffs of clean air or EC aerosols, followed by 24 hrs of exposure to pseudoparticles. Exposure was at the ALI in a Cultex® system. All graphs are plotted as the mean ± standard deviation of three independent experiments. Box-Cox transformed data were analyzed using a one-way ANOVA followed by Tukey’s test to compare means. Black * = significantly different than the control in **E** or the exposure control in **G and H.** Color * = significantly different groups. *p < 0.05, **p < 0.01, ***p < 0.001, ****p < 0.0001.

After 24 hours of submerged treatment, viral pseudoparticles were added to the cell cultures for 24 hours, then fluorescence was examined using microscopy and flow cytometry (**Figure 4B**). Infection increased significantly in cells treated with authentic JUUL™ “Virginia Tobacco” fluid (**Figure 4 C, E**). Infection increased slightly, but not significantly, in PV/VG alone. Nicotine significantly increased infection dose dependently, and infection was slightly higher when nicotine was combined with PG/VG, rather than used alone (**Figure 4C, E**).

Similar results were obtained with BEAS-2B cells exposed to nicotine-containing aerosols at the ALI in a cloud chamber (**Figure 4G**). Infection was significantly elevated in the group exposed to 0.3 mg/mL of nicotine and was close to significance (p = 0.06) in the group exposed to the lower concentration of nicotine (**Figure 4G**). ALI exposure to various aerosols in the Cultex® system significantly increased infections in all treatment groups with the effect being strongest in the groups that had nicotine (**Figure 4H**). The increase in infection was significant in both the JUUL™ aerosols and the high nicotine plus PG/VG group; however, the increase in the JUUL™ group was significantly lower than in the nicotine PG/VG group. These increases in infection are likely due to higher ACE2 expression (**Figure 1**) and elevated TMPRSS2 activities (**Figure 2–3**) in the nicotine-treated BEAS-2B cells.

## Discussion

Our results, summarized in **Figure 5**, show the effects of exposure protocol (submerged vs ALI), heated vs unheated aerosols, and authentic JUUL™ aerosol versus individual aerosol chemicals on the machinery involved in SARS-CoV-2 infection. These results support the conclusion that JUUL™ “Virginia Tobacco” e-liquids and aerosols can increase ACE2 levels, TMPRSS2 activity, and infection of BEAS-2B bronchial epithelial cells by SARS-COV-2 pseudoparticles. By examining different exposure protocols (submerged vs ALI) and chemical variables (fluids, aerosols, PG/VG, and nicotine), we show that the effects of ECs on viral infection depend on both the context in which exposures occur (submerged vs ALI) and the chemical formulations of the aerosols *per se*. In submerged cultures and cloud chamber exposures, increases in ACE2 levels and infection can be attributed mainly to nicotine. In the Cultex® system, these increases were associated with both nicotine and PG/VG, perhaps due to formation of reaction products (e.g., aldehydes, ketones and alcohols) from the solvents during heating [44, 53, 54, 55, 56]. Metals (e.g., zinc, copper, iron, lead, nickel, tin) are also added to aerosols during heating [46, 48, 49], and these could also influence the outcome. While both the levels and activity of TMPRSS2 increased in submerged cultures, only activity increased in ALI exposures. Increases in TMPRSS2 activity were associated mainly with nicotine, while PG/VG produced a small but significant decrease in the Cultex® system. While JUUL™ aerosols increased infection significantly, this increase was not as great as in the group containing only PG/VG/nicotine in concentrations equivalent to those in JUUL™, showing both that JUUL™ e-liquid decreased the effects of nicotine and the importance of testing authentic EC aerosols, such as JUUL, which modulated the effectiveness of nicotine. For most endpoints, both submerged and ALI exposures led to similar conclusions, but the ALI provides data on an exposure similar to that an EC user receives. Moreover, the Cultex® exposure enables testing of authentic EC aerosols and is the best protocol for evaluating the human response.

**Figure 5:**
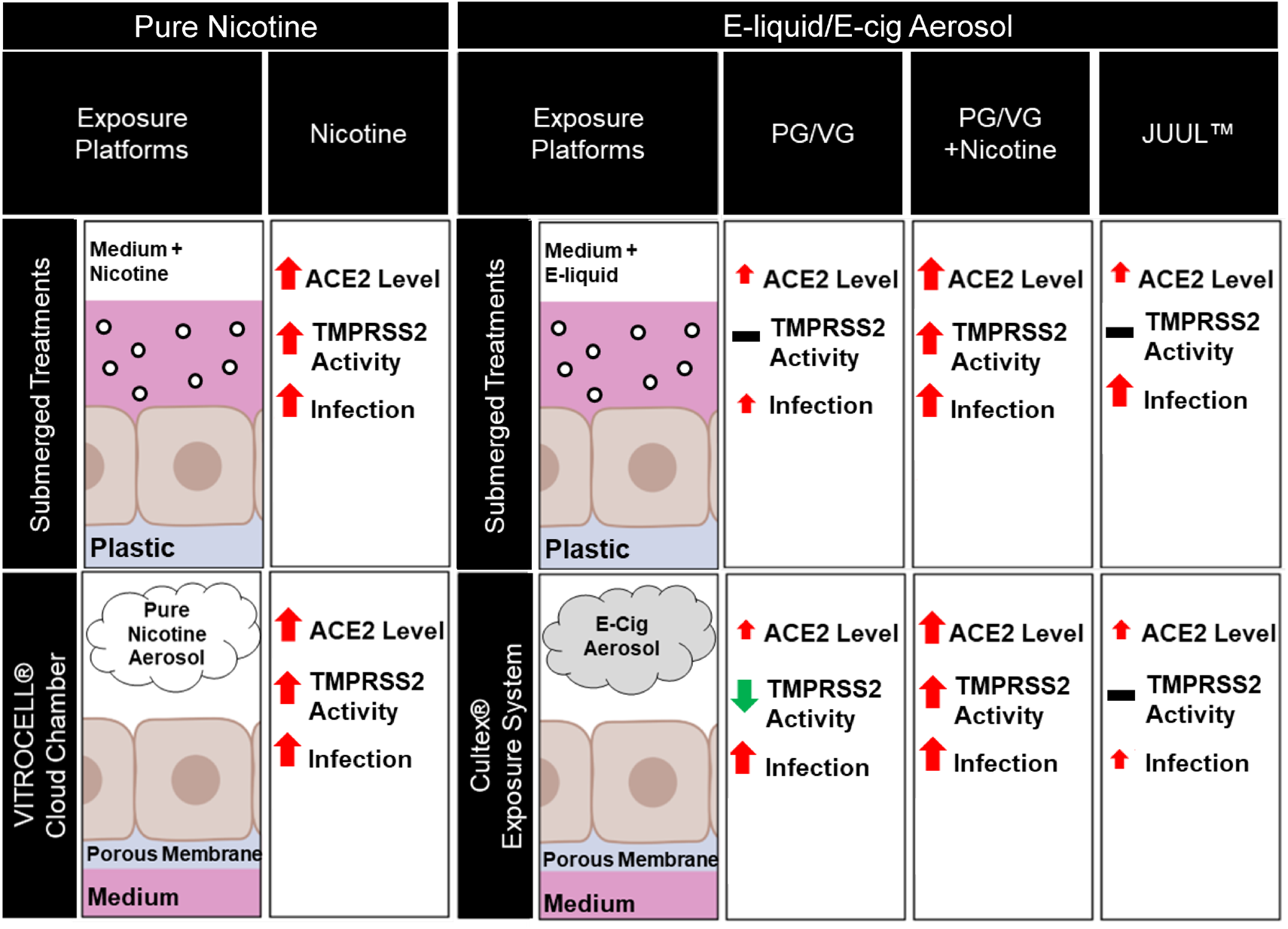
Relationship Between JUUL™ E-liquid, Nicotine, and PG/VG Exposure and ACE2, TMPRSS2, and Viral Pseudoparticle Infection. Diagram summarizing our major findings in BEAS-2B cells following exposure to authentic JUUL™, mixture (PG/VG plus nicotine), and individual chemicals (nicotine and PG/VG solvent) in three different exposure models: submerged treatment (top), air-liquid interface (ALI) exposure in the cloud chamber (bottom left), and ALI exposure of authentic EC aerosols in a Cultex® system (bottom right). ACE2 levels increased with all treatments in each exposure platform. TMPRSS2 activity varied with the platform and treatment. Most treatments increased TMPRSS2 activity, three treatments had no effect (PG/VG in submerged and PG/VG and JUUL in the Cultex exposures), and one treatment decreased activity (PG/VG in the Cultex). Infection was increased in all treatments/exposures. Size of the arrow indicates relative change from the control.

Table 1 summarizes the nicotine concentrations in our three exposure platforms (fluid concentrations in submerged exposures and concentrations measured in insert culture media after ALI exposures) and compares them to estimated nicotine concentrations in human alveolar lining fluid after smoking a cigarette [57, 59]. We adjusted exposures in the two ALI platforms to produce similar final concentrations of nicotine in the insert fluids after exposure. As seen in Table 1, the nicotine concentrations in both the Cultex® and cloud chamber inserts were similar and were within the range reported in the alveolar lining fluid of smokers after smoking one cigarette [57, 58], supporting the conclusion that our ALI exposures were reasonable and representative of those received in vivo by humans during vaping. A much higher starting concentration, which is similar to concentrations found in actual EC products, was used in the Cultex® system as the aerosol is diluted before entering the exposure chamber to ensure uniform distribution. In contrast, undiluted aerosol can be used in the cloud chamber, where it disperses uniformly and hence a lower starting concentration and puff number was used. These data demonstrate that chemical exposures were similar between the cloud chamber and Cultex® system and that exposures in the ALI systems, based on nicotine concentration, were within the range a tobacco product user would have in their alveolar lining fluid.

**Table 1:** Nicotine concentrations across the in vitro platforms compared to nicotine concentrations in alveolar lining fluid of human smokers^1^

| Platform | Concentration in submerged culture fluid and aerosolized fluid | Concentration in culture fluid and insert fluid | Concentration in alveolar lining fluid of smokers after 1 cigarette<sup>1</sup> |
| --- | --- | --- | --- |
| Submerged Culture | 30 and 300 µg/mL | 30 and 300 µg/mL | 1 -10 µg/mL |
| Cloud Chamber ALI Exposure | 30 and 300 µg/mL | 1.3 µg/mL | 1 -10 µg/mL |
| Cultex® ALI Exposure | 6 and 60 mg/mL | 1.5 µg/mL Authentic JUUL™<br>1.8 µg/mL Lab-made Nicotine | 1 -10 µg/mL |

<sup>1</sup> Ranges are taken from references in the [57, 58].

The effective concentrations and length of exposure to nicotine differed across exposure platforms. Nicotine increased SARS-CoV-2 pseudoparticle infection of BEAS-2B cells in all exposure systems; however, in submerged culture, exposures were continuous over 24 hours, while in the ALI systems, exposures were intermittent and short in duration, suggesting cells are more sensitive when exposed at the ALI. Our data are in agreement with other studies showing that zinc nanoparticle exposure of A549 cells was more likely to produce an effect in ALI exposures than in submerged cultures [60] and that gaseous exposure to aldehydes at the ALI caused significantly higher levels of IL-8 secretion than submerged culture exposures [61]. The apparent increase in cell sensitivity at the ALI may come about because nicotine or other chemicals in the EC aerosols were not diluted by the medium and interact directly with the surface of cells in the absence of culture medium. Because the effects of nicotine on infection are concentration dependent, results of both the ALI and submerged exposures are likely to vary with EC products, which are available in a broad range of nicotine concentrations [25, 62]. Our in vitro models support the conclusion that cells are more responsive to nicotine at the ALI than when submerged and that nicotine dose dependently increases ACE2 levels, TMPRSS2 activity, and SARS-CoV-2 pseudoparticle infection.

One of our goals was to examine the effects of individual chemicals in JUUL™ aerosols on ACE2, TMPRSS2, and infection. JUUL™ “Virginia Tobacco” was chosen because its concentrations of nicotine and PG/VG are known, and it has relatively few flavor chemicals, which are used at low concentrations [35]. Our experiments compared authentic JUUL™ “Virginia Tobacco” aerosols to aerosols made with fluids containing PG/VG, nicotine, and/or PG/VG/nicotine mixtures using chemical concentrations that replicated those in JUUL™ “Virginia Tobacco” fluid. JUUL™ “Virginia Tobacco” aerosol increased the levels of ACE2 in BEAS-2B cells across all platforms, in agreement with and extending prior work showing that submerged treatment of BEAS-2B cells with Watermelon-flavored refill fluid increased ACE2 transcripts in BEAS-2B cells [34]. JUUL™ fluids also increased ACE2 enzymatic activity in vitro [20] although ACE2 serves as a spike protein receptor and its activity is not required for infection [63]. Our data isolated these increases to nicotine and PG/VG in the Cultex® system. Our data further showed that: (1) nicotine increased ACE2 levels dose dependently, (2) the nicotine concentration in JUUL™ “Virginia Tobacco” was sufficient to significantly elevate ACE2, and (3) the concentrations of nicotine reaching insert media in both the cloud chamber and Cultex® system were within the range reported in smokers alveolar lining fluid [57, 58]. Based on our data and other publications [32, 64, 65], there is a consensus that nicotine increases the ACE2 receptor in human bronchial epithelial cells.

Most prior SARS-CoV-2 studies have focused on ACE2 expression [20, 30, 31, 33, 34], and did not examine the effects of EC chemicals on TMPRSS2 concentration and activity. Increased TMPRSS2 activity would promote spike cleavage at the S2’ site, internally to the S2 subunit, and fully activate the viral fusion process [12, 66]. Our data show that PG/VG, nicotine, and JUUL™ aerosols had different effects on TMPRSS2 activity. In most cases, TMPRSS2 activities were significantly increased by nicotine. An interesting exception was the lack of effect of JUUL™ “Virginia Tobacco” aerosols on TMPRSS2 activity, even though JUUL™ aerosol had a nicotine concentration similar to that in the group with 60 mg/mL of pure nicotine in PG/VG. This again shows the importance of examining authentic EC aerosols as the effects of nicotine may be modulated in complex aerosols containing mixtures of chemicals.

In summary, our data show that JUUL™ “Virginia Tobacco” aerosols modulated the infection machinery in BEAS-2B cells in a manner that increased infection by SARS-CoV-2 viral pseudoparticles. However, nicotine in PG/VG was more effective than nicotine in JUUL™ aerosol, showing that chemicals in authentic JUUL™ aerosols can reduce nicotine’s effect. In ALI exposures, increases in infectivity were generally correlated with nicotine and/or PG/VG induced increases in ACE2 concentrations and increases in TMPRRS2 activity. Results were generally similar across the three exposure platforms, but exposures in the ALI systems were brief and intermittent and involved lower concentrations of nicotine than in submerged cultures, which received 24 hours of continuous exposure. Given the highly variable nature of EC devices and their fluids [67], it is probable that infection by SARS-CoV-2 is influenced by the product, user topography, and the e-liquid used for aerosol generation. Our data identify chemicals in EC liquids and aerosols that affect SARS-CoV-2 infection, show that JUUL™ EC aerosols can modulate infection, and identify nicotine and PG/VG as chemicals that can increase SARS-CoV-2 infection of human bronchial epithelium. Given the number of factors that can influence infection, it is not surprising that prior studies on tobacco products, mainly conventional cigarettes, have sometimes differed in their conclusions regarding the relationship between smoking/vaping and COVID-19 [5, 6, 7, 13, 14, 15, 16, 17, 18, 68]. EC products are highly variable and have nicotine concentrations ranging from 0 to 60 mg/mL [21, 22, 61, 69]; operate at different powers, have different atomizer designs [70], and have varying chemical mixtures in their fluids, all of which can affect SARS-CoV-2 infection and severity differently. Based on our BEAS-2B data, it is probable that some EC products deliver sufficient nicotine to increase infection, while those that are nicotine-free or have low concentrations of nicotine may not influence SARS-CoV-2 infection.

Our data support the conclusion that understanding the relationship between EC use and COVID-19 will be challenging due to the many variables related to vaping. Variations in EC design, power, user topography, and e-liquid ingredients are some of the factors that can influence SAR-CoV-2 infection. These variables notwithstanding, our data can be valuable in improving the safety of EC. For example, since nicotine increased infection dose dependently, capping the allowable concentration of nicotine in e-liquids could reduce the likelihood of SARS-CoV-2 infection in EC users.

### Limitations of the Study

We examined the effect of EC aerosols on BEAS-2B cells. In the future, it will be important to extend these studies to a 3D organotypic model of the respiratory epithelium that closely resembles *in-vivo* exposure in humans. Our study is limited to one SARS-CoV-2 variant and could in the future be extended to new variants as they arise. Our work included JUUL™ “Virginia Tobacco”, which operates with a relatively low voltage and power. Variations in power can affect formation of reaction products, which may in turn alter infectability of cells. Finally, our exposures were acute and could be extended to chronic exposure in the future.

## Abbreviations

EC: electronic cigarettes
PG: propylene glycol
VG: vegetable glycerin
ALI: air-liquid- interface
ACE2: angiotensin converting enzyme 2
TMPRSS2: transmembrane serine protease 2
COVID-19: corona virus disease-2019
SARS-CoV-2: severe acute respiratory syndrome coronavirus 2
MOI: multiplicity of infection

## Acknowledgements

We thank Dr. James Pankow, Dr. Wentai Lou, and Kevin McWhirter from Portland State University for quantifying the deposition of nicotine in the ALI exposure systems. We would like to acknowledge Jack Ono and Claudia Osuna for assisting in viral pseudoparticle production. The Diagram in Figure 5 was created with BioRender.com. We thank the UCR stem cell core for providing the instrumentation for flow cytometry.

## Authors’ contributions

Project administration and funding acquisition, P.T.; Conceptualization, R. P. and P.T.; Investigation R.P.; Sample preparation, data collection, and data processing were done by R.P., M.W., A.S., and T.M.; Data Interpretation was done by R.P. and P.T.; Writing the original draft was done by R.P. and P.T. Reviewing and editing was done by RP, PT and TM.

## Availability of data and materials

The datasets analyzed in this are available from the corresponding author upon request.

## Ethics approval and consent to participate

Not applicable.

## Consent for publication

Not applicable.

